# Climatic determinants of plant phenology in vernal pool habitats

**DOI:** 10.1101/2024.07.25.605166

**Authors:** Brandon Thomas Hendrickson, Aubrie Heckel, Robert Martin, Jason Sexton

**Affiliations:** Department of Biology, University of Louisiana at Lafayette; Department of Life and Environmental Sciences, University of California, Merced

**Keywords:** phenology, ephemeral wetlands, growing degree hours, climate change, endemic, citizen science

## Abstract

Vernal pool plants are small, colorful, and specialized to both desiccated and inundated conditions that distinguish the ephemeral wetlands in which they grow. These species germinate rapidly in response to the first rain and grow quickly to take advantage of available water supplies. The floral phenology of vernal pool plant species is little understood despite being a crucial developmental stage for producing seeds and determining population growth rates. The current study focuses on two vernal pool plants, *Limnanthes douglasii* ssp. *rosea* (meadowfoam), a vernal pool specialist, and *Trifolium variegatum* (whitetip clover), a generalist vernal pool associate, and characterizes their phenology in response to interannual climate variation. We recorded phenology and climate data over seven years during a period of highly variable precipitation and temperature patterns, which serve as a robust dataset for quantifying the relationship of floral phenology with various climatic factors. We found that warmer and drier environmental conditions occurring during early growth periods were strongly associated with advanced floral phenology later in the life cycle for both species. Over the seven-year dataset, which was increasingly warm and dry, phenology advanced by 4.7 days per year for meadowfoam and 5.6 days per year for whitetip clover, respectively. The floral duration of the habitat specialist was influenced by microtopographic features of vernal pools, whereas no such patterns were observed for the habitat generalist. Finally, warmer and drier conditions were associated with reduced occupancy rates of both focal species within vernal pools. To our knowledge, this is the first study quantifying the relationship between vernal pool floral phenology and climate, offering insights into how phenology may shift in response to modern climate change.

## INTRODUCTION

The onset of flowering acts as a fingerprint of climatic variation in an ecosystem (Parmesan & Yohe, 2003; Fatima et al., 2020; Menzel et al., 2020) and dictates the window of time during which symbiont associates can capitalize on plant resources (Benaldi et al., 2013; Polsedovich et al., 2018). Vernal pool habitats, ephemeral wetlands that are now rare and endangered in California, harbor many endemic plant species (Holland, 1976; Stone, 1990; Witham, 1998) adapted to sharp transitions from extreme wet to extreme dry conditions (Keeley & Zedler, 1998). Vernal pools annually fill with water during the winter and dry over the spring season, offering a unique system for investigating how climate in the winter and spring independently affect growth and adult flowering phenology. The phenological dynamics of specialist species and more generalist species growing in vernal pools have not been adequately studied in the context of climate variation. Consequently, neither the abiotic cues that trigger flowering nor the degree to which flowering time, length, and termination change in response to climate is currently understood in vernal pool habitats. We aimed to quantify the relationship between seasonal climate conditions and interannual phenological patterns of two vernal pool plant species (one specialized, the other more generalist) over a seven-year study to better understand the susceptibility of vernal pool habitats to present and future climate change. We conducted this research in a community science context, involving undergraduate student volunteers from the University of California, Merced, every spring in the collection of phenology data.

The environmental factors driving phenological transitions remain unknown in natural vernal pool habitats. Controlled experiments in constructed mesocosms have found significant associations between flowering onset and termination with inundation length; however, the direction of the response was variable among plant species (Collinge et al., 2013). These varying floral responses suggest that vernal pool species exhibit unique phenological responses to hydroperiod or other previously unexplored environmental factors, such as temperature. Although hydroperiod can predict germination and plant development timing of vernal pool species (Collinge et al., 2013), flowering time networks studied in model plant species have revealed multiple exogenous factors, such as temperature and photoperiod (Amasino, 2010). Additionally, plant species with wider niche breadths, occurring in but not restricted to vernal pools, may be less influenced by changes in inundation than vernal pool specialists. Studying both widespread and specialist species in a natural setting can test how phenology of vernal pool plants is influenced by a diversity of climatic factors and elucidate whether these species are susceptible to future climate change (Shay et al. 2021).

Climate change is manifesting in several ways, including increasing temperatures, drought, and weather variability (Bauder, 2005; Brooks, 2009; Dettinger & Cayan, 1995). Global patterns of shifting floral initiation in response to warming temperatures and lower precipitation have been recorded on six of the seven continents and in many biomes (Post et al., 2018; Karouba et al., 2018). However, phenological studies have not yet included vernal pool habitats, which are ideal for understanding ecological patterns in ephemeral environments (e.g., Ruiz-Ramos et al., 2022). The degree to which climate is changing depends on the season; fall minimum and maximum temperatures are rising more rapidly than spring temperatures compared to historical averages in the Central Valley of California (He et al., 2018). Given that vernal pool species emerge in fall and bloom in spring, it is important to investigate which season has a larger impact on bloom start and end dates. Additionally, there is currently no clear pattern of how flower duration will change in response to warmer and drier conditions (Li et al., 2020; Puchalka, 2022).

Agricultural expansion has destroyed 95% of the original vernal pool habitat native to California (Ruiz-Ramos et al., 2022). Habitats with species loss caused by human encroachment, invasion, and fragmentation, such as California’s vernal pools, are closer to complete ecosystem collapse than those that evolved with human activity (Prim & Raven, 2000). Consequently, many vernal pool organisms are endangered and continuously threatened by invasion and human encroachment, elevating the need for understanding the role climate change may play in further disrupting this already fragile habitat.

We studied several potential abiotic and seasonal drivers of vernal pool floral phenology in situ and subsequently determined the interannual patterns of flower onset, termination, and duration in response to variable temperature and precipitation. During the first half of the growing season, vernal pool plants must germinate and establish when pools are inundated and daily temperatures are metabolically restrictive for further growth. Plants then transition to vegetative growth and reproduction typically during the latter half of the growing season as water recedes and air temperatures rise. By dividing the winter season into two periods (i.e., early winter and late winter seasons), we examined the effects of climate on early and late vernal pool phases independently. We ask three questions: 1. What climatic factors best explain vernal pool plant phenology? 2. Is the phenology of vernal pool specialist and generalist plant species responding to climatic variation similarly? And 3. How does floral duration change in response to climatic variation and pool topography?

## METHODS

### 2.1 Study System

The study was conducted in Merced County within the Central Valley of California on the University of California, Merced Vernal Pool & Grassland Reserve (MVPGR). Cattle grazing is used as a management tool here to control invasive plants and preserve other ecosystem functions (Barry 1995). Merced falls between the hot-summer Mediterranean and semi-arid steppe climate regions of California as described by the Koppen climate types (Kesseli 1942). Interannual climate variability is high for this climate region, with frequent droughts and flooding. Mediterranean climates are marked by extreme seasonal cycles, from cold, inclement winters to hot, desiccating summers that have selected for a unique assemblage of plants with rapid life cycles. The vernal pools are fed by rainfall during the winter, the frequency, amount, and timing of which, combined with the carrying capacity of a pool, dictates the length of the growing season (Zedler 2003). We studied three observation pools within the MVPGR differing in several environmental aspects, including clay content, depth, diameter, and soil type (Figure 1).

**Figure 1:**
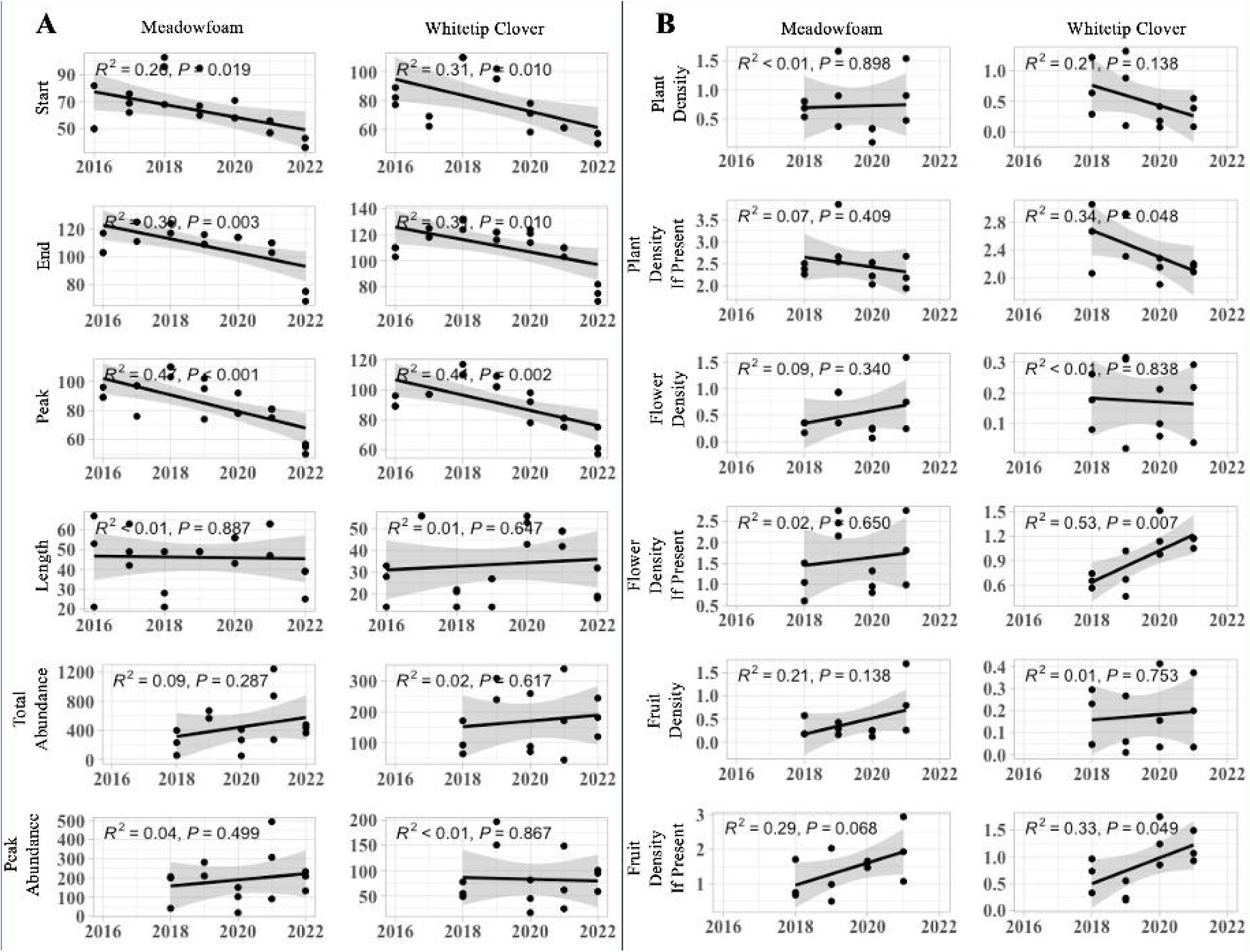
(A) Map of the Merced Vernal Pool and Grassland Reserve (MVPGR) with the locations of three observation pools investigated from 2016-2022. Adapted from Swarth 2021. Included are is the location of the Merced Vernal Pool Megacomplex denoted by a red star on the California map (B) A cohort of undergraduate citizen scientists recording the density of plants, flowers, and seeds of meadowfoam and whitetip clover. (C) A vernal pool during peak bloom with white meadowfoam flowers in the center and whitetip clover along the edges. Photos of the two focal species, (D) whitetip clover (*Trifolium variegatum)* and (E) meadowfoam (*Limnanthes douglasii* ssp. *rosea*). Photography credits to (C) Monique Kolster, and (D) Barry Rice.

Vernal pools are depressions of approximately a half meter in depth underlain by an impermeable soil horizon (Holland & Jain 1988; Boone et al., 2006), either clay or rock, that cyclically flood during the wet season, gradually dry throughout the spring, and then dry during the summer (Keeley & Zedler 1998; Solomeshsch et al., 2007). Few plant species can tolerate such extremes of water availability, and consequently, the flora is composed of many endemics that inhabit the deeper portions of vernal pools (Hanes & Stromberg 1998; Keeley & Zedler 1998) and other more widespread species growing along the pool margins and surrounding grasslands. Predictably sharp transitions from inundated to desiccated that characterize vernal pools can be divided into two biologically informative temporal phases: (1) the aquatic phase when plants quickly germinate following the first rain and grow slowly as diminutive rosettes while submerged, and (2) the terrestrial phase that commences as water recedes and vernal pool plants rapidly complete their life cycle while soils are saturated (Keeley & Zedler 1998; Pollak & Kan 1998; Bauder 2000; Gerhardt & Collinge 2003). Much knowledge of vernal pool plant phenology has been acquired using this heuristic, such as correlating hydroperiod with seedling emergence (Bliss & Zedler 1998; Collinge et al., 2013), community composition (Barbour et al., 2003; Deil 2005; Solomeshch et al., 2007; Gosejohan et al., 2017), phylogenetic diversity (Hendrickson 2024), and plant traits (Kraft et al., 2014)

The two focal species in this study were whitetip clover, *Trifolium variegatum*, and rosey meadowfoam, *Limnanthes douglassii* ssp. *rosea*. Meadowfoam is an annual, California endemic species and a vernal pool habitat specialist. The cream-colored flowers are often observed in bloom between early March to late April at the bottom of vernal pools (Hickman 1993). The range of meadowfoam is restricted to hardpan and claypan pools in the southeast of San Joaquin Valley (Keeler-Wolf et al., 1998). This contrasts with the more generalist species, whitetip clover, which inhabits a wide range of habitats from Alaska to Baja California (Čelakovský 1874; Hickman 1993) and can occupy various elevations within a vernal pool (Bliss & Zedler 1998). Whitetip clover is known to flower from March to July across its range (Calflora, https://www.calflora.org), however, to our knowledge, its phenological patterns have not yet been extensively described within vernal pool habitats. We monitored the phenology and demographics of both species for seven years, including within year and between year precipitation and temperature patterns, to relate phenological variation with climate variation.

### 2.2 Characterizing the Growing Season

Aquatic or wet-season biological activity in vernal pools is largely concentrated from October to May (Zedler 2003). The consequence of a truncated growing season in vernal pools is that annual averages of precipitation and temperature are often uninformative of growth patterns of vernal pool plant communities (Javornick & Collinge 2016); thus, studies of community composition and phenology have focused on late fall, winter, and early spring weather patterns. Prior studies of California vernal pool plants have shown important transitions from germination, survival, growth, and reproduction that parallel seasonal shifts of water volume (Keeley & Zedler 1998; Collinge et al., 2013; Gosejohan et al., 2016). Given this important distinction between growth stages driven by inundation and temperature, we followed Javornik & Collinge (2016) to differentiate between early winter (October – December) and late winter (January – March) seasons.

### 2.3 Phenology Census

Weekly visits to 3 study vernal pools on the Merced Vernal Pool and Grasslands Reserve started in late January to visually inspect for the presence of meadowfoam and whitetip clover flowers. Once flowers were observed, a team of undergraduate volunteer researchers laid out north-south and east-west transects spanning the length of each pool (Figure 1). Each year we monitored plant abundance and phenological transitions each week until May 30^th^ or until flowers senesced, whichever occurred first. Quadrats (10 cm × 10 cm) were placed every 20 cm alongside each transect and the number of plants, flowers, and fruits within each quadrat was counted.

Meadowfoam has tendril-like branches that creep upon the pool surface that can occasionally extend several inches from the base of the plant. We counted plants of both species only when rooted within the quadrat. Flowers occurring within a quadrat were counted once anthesis had occurred. The fruiting phenophase of meadowfoam was determined when sexual organs fully degenerated and nutlets became clearly visible. Whitetip clover produces fruits (legumes) covered by dried sepals; however, the flower is visibly dry and the petals brown when legumes are being produced, which served as the indicator of the plant’s transition to fruiting. The first flowering date for each focal species was defined as the day when any flowering plant in a pool was first observed, and the end date was defined as the date when no more flowers were visible in a pool. Two additional statistics were derived from the total number of flowers within a pool: cumulative abundance of flowers over a growing season and maximum abundance of flowers within a growing season. Additionally, the date of peak abundance of flowers was recorded.

### 2.4 Characterizing Temperature and Precipitation

Regional precipitation and temperature were derived from measures at the Merced Field Station (37.314, -120.387), managed by the California Irrigation and Management Information System approximately 10 km from the study site. We downloaded daily measurements of several precipitation and temperature metrics from October 1^st^, 2015, to May 30^th^, 2022: accumulated precipitation, minimum temperature, maximum temperature, and mean temperature. For each year and month, we calculated the cumulative precipitation, average maximum temperature, average minimum temperature, and average mean temperature. We also calculated the same metrics for early and late winter periods. To determine how our seven-year study period compared to the average climate patterns of Merced during the past few decades, we also collated data from October 1^st^, 1999 (the first year the Merced climate station became active), to May 30^th^, 2022.

Growing degree hours were calculated as the summation of hours in a day, month or year that exceed a temperature threshold. We chose a base temperature of 9.44° C as the threshold following the suggestion by Javornik & Collinge (2016) that vegetation in vernal pool environments commence growth when mean air temperature reaches 9.44° C. We downloaded hourly temperature data from October 1^st^, 2015, to May 30^th^, 2022 and calculated the accumulated growing degree hours for early winter (October – December) and late winter (January – March) growing seasons. Given that GDH is dependent upon a thermal threshold, diurnal temperature variability is likely to influence the accumulation of GDH. To evaluate the relationship between GDH and daily temperature variation, we calculated the daily temperature range and used multiple regression with temperature variability and maximum daily temperature as predictor variables of GDH.

Climate moisture index (CMI) was calculated for each day to evaluate drought conditions in the vernal pools using the following equation:

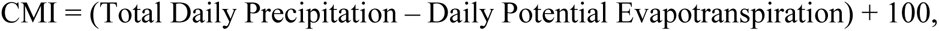

with daily potential evapotranspiration calculated using the Penman-Monteith equation (Monteith 1965). Average daily CMI for early winter and late winter growing seasons were calculated each year.

Given that the Merced climate station was 10 km from the study site, we tested whether the Merced climate station data were representative of regional precipitation patterns. To do this, we compared the accumulated rainfall recorded at four nearby CIMIS climate stations: Denair II, Fresno State, Oakdale, and Orange Cove. The closest and farthest climate stations to Merced were 35 and 128 km away, respectively. We performed ANOVA with the car package and Dunnett’s test using the desctools package (Signorell 2023) in R (v.4.2.0, R Core Team 2021) to determine if Merced was significantly different than the 4 adjacent climate stations.

Finally, we quantified the interannual trends of precipitation, GDH, and mean temperature across the study period using linear regression of these variables with year. An ANOVA was used to test if early winter climate was significantly different than late winter climate.

### 2.5 Pool Topographic Characteristics

To determine if fine scale microtopographic variation affects the phenological patterns of our focal species, we measured the diameter, depth, and soil texture of each pool (Table 1). Pools with a large diameter and depth can retain water for a longer period than smaller pools. The length and width of each pool was measured using a transect that spanned the entire pool bottom and approximately 2 meters into the grassland. The depth of each pool was measured in January of 2018 – 2020 using a meter stick at the center of each pool.

**Table 1.**
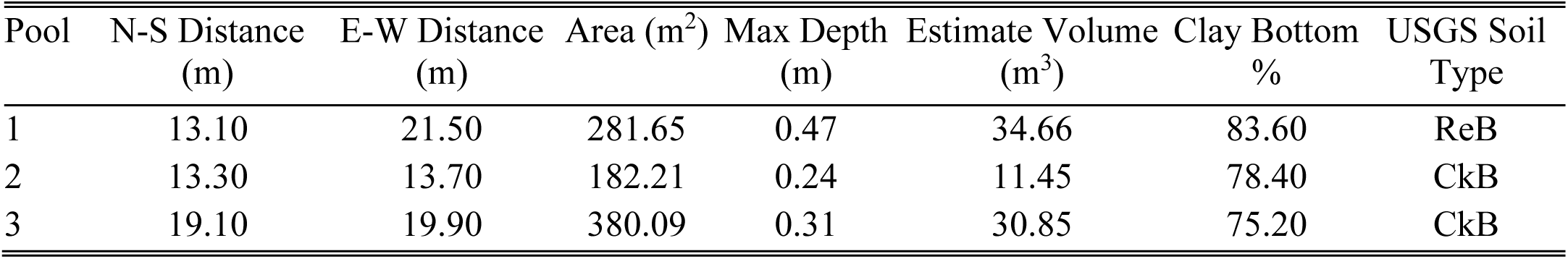
Dimensional and edaphic characteristics of three observation pools on the Merced Vernal Pool and Grassland Reserve. North-South (N-S) and East-West (E-W) distances were measured using a transect spanning the pool bottom and 2 meters into the pool upland. Pool area was calculated as the elliptical area of the pool. Max depth was the height of water in the center of a pool during the wettest year and while the pool was inundated. Estimated volume was calculated by using the maximum depth of water and the elliptical area of the pool to determine the hemispheric volume. Clay percentage is the average clay content found throughout the bottom of each pool. Soil types described by USGS soil survey are Redding gravelly loam with 0 to 8 percent slopes (ReB) and Corning gravelly sandy loam with 0 to 8 percent slopes (CkB).

We determined the soil type of the pools using a land survey dataset produced by USGS (https://websoilsurvey.nrcs.usda.gov/app). Additionally, we estimated soil texture gradients within pools by sampling ca. 50g of topsoil every meter along a north-south and east-west transect during the dry season. Five soil replicates per sampling point were measured and averaged. We used an assessment-by-touch method to estimate clay, sand, and silt content (Thien 1979).

### 2.6 Analyses of Phenology Responses to Climate

We tested for the normality of each phenological variable using the Anderson-Darling statistic with the nortest R package (Zeileis & Hothorn 2002). The assumption of normality was met for six out of eight dependent variables; meadowfoam end date and whitetip clover end date were non-normal. To determine if nonparametric tests were necessary, we first log transformed the non-normally distributed variables. We then ran 5 sets of multiple regressions with different predictor variables on both transformed and untransformed data in base R: (1) GDH and precipitation, (2) GDH and CMI, (3) Mean temperature and precipitation, (4) Minimum temperature and precipitation, (5) Maximum temperature and precipitation. We compared the AIC value of models using transformed data and untransformed phenological measurements as in Javornik and Collinge (2016). Untransformed variables performed better in all model scenarios. Thus, we chose to report and assess the results obtained using untransformed data.

To test for effects of climate variables on phenological responses, we ran linear multiple regression models in a stepwise fashion for each phenological variable against all climate variables using the *lm* function in base R, removing nonsignificant predictor variables with the highest P value until every value was statistically significant at the alpha = 0.05 level. Furthermore, AIC was calculated for each model to determine which performed better. Models with the lowest AIC score were reported. All model residuals were tested for patterns of heteroscedasticity using a Breusch-Pagan test using the lmtest package (Bates et al., 2015) in R.

### 2.7 Analyses of Phenology Responses to Pool Topography

In order to test the effect of pool topography on phenology, we ran linear regression models of flowering start date, end date, and length of flowering with “distance from edge” as the predictor variable. Distance from edge is the distance a given quadrat is from the nearest edge in the pool. Linear regression models were performed using the *lm* package in R.

### 2.8 Quantifying Impacts of Shifting Phenology on Abundance

We used multiple regression models to quantify the impact warmer climates and shifting flowering times have on plant density. Total winter precipitation, total winter GDH, and flowering start and end time were independent variables. Flower density, fruit density, and plant density were treated as response variables. To distinguish between density estimates across a pool, versus density across occupied plots, we calculated two average values. The first average, which we call “pool density,” includes both occupied and unoccupied quadrats (quadrats with zero values). The second average, which we call “plot density,” only includes quadrats with at least one plant present. Finally, we tested if predictor variables were significantly associated with “occupancy” rates within a pool, calculated as the number of quadrats with at least one plant present divided by the total number of quadrats in a pool. We calculated the average occupancy for each year of a given pool, and we standardized the average value by pool size. Multiple regressions were performed in base R.

## RESULTS

### 3.1 Climate Patterns

The Merced climate station represented a viable regional proxy for climate patterns at the three study pools. An ANOVA test showed no significant differences among stations regarding accumulated rainfall in either the late or early winter seasons over the course of the study period (Early Precipitation: F = 0.1537, P = 0.9598; Late Precipitation: F = 0.1263, P = 0.9718). Additionally, Dunnett’s test with Merced as the control showed that early and late winter precipitation patterns in Merced were not significantly different from other climate stations (Supplemental Figure 1).

Winter climate showed significant interannual variability during the study period (Figure 2). Between 2016-2022, early winter precipitation fluctuated substantially from a low of 38.8 mm in 2018 to a high of 148.9 mm in 2022 (Table 2), whereas late winter precipitation had a low of 34.6 mm in 2022 and a high of 325.6 mm in 2017. Linear regressions of annual precipitation over the 7-year study period in the early winter and late winter yielded a significantly negative correlation only for late winter rainfall (p = 0.048; ß = -37.34; R2 = 0.577, F = 6.83). The range of accumulated growing degree hours (GDH) in early winter was 425, with a minimum of 892 in 2018 and a maximum of 1317 in 2022 (Table 2). Accumulated GDH in late winter ranged from 973 in 2021 to 1213 in 2016. From 2016-2022, growing degree hours in early winter were not correlated with year (p = 0.514; ß = 17.93; R2 = 0.090; F = 0.4924), whereas late winter GDH showed a significantly negative correlation with year (p = 0.011; ß = -36.64; R2 = 0.755, F = 15.4). Daily mean temperature in late winter was trending colder (p = 0.119; ß = -0.144; R2 = 0.41; F = 3.53) whereas daily mean temperature in the early winter was trending warmer (p = 0.435; ß = 0.054; R2 = 0.1257; F = 0.72), though neither trend was significant. Minimum temperatures in late winter were significantly colder each year (p = 0.0008; ß = -0.48; R2 = 0.91, F = 50.99) and maximum temperatures in late winter trended warmer each year (p = 0.0824; ß = 0.32; R2 = 0.48, F = 4.7), though this relationship was not significant. Diurnal temperature variability was positively associated with year in the late winter (p = 0.00365; ß = 0.86; R2 = 0.84, F = 26.4). The CMI in the late winter was falling during the study period (p = 0.0289; ß = - 0.86; R2 = 0.84, F = 26.4), indicating more drought-like conditions.

**Figure 2:**
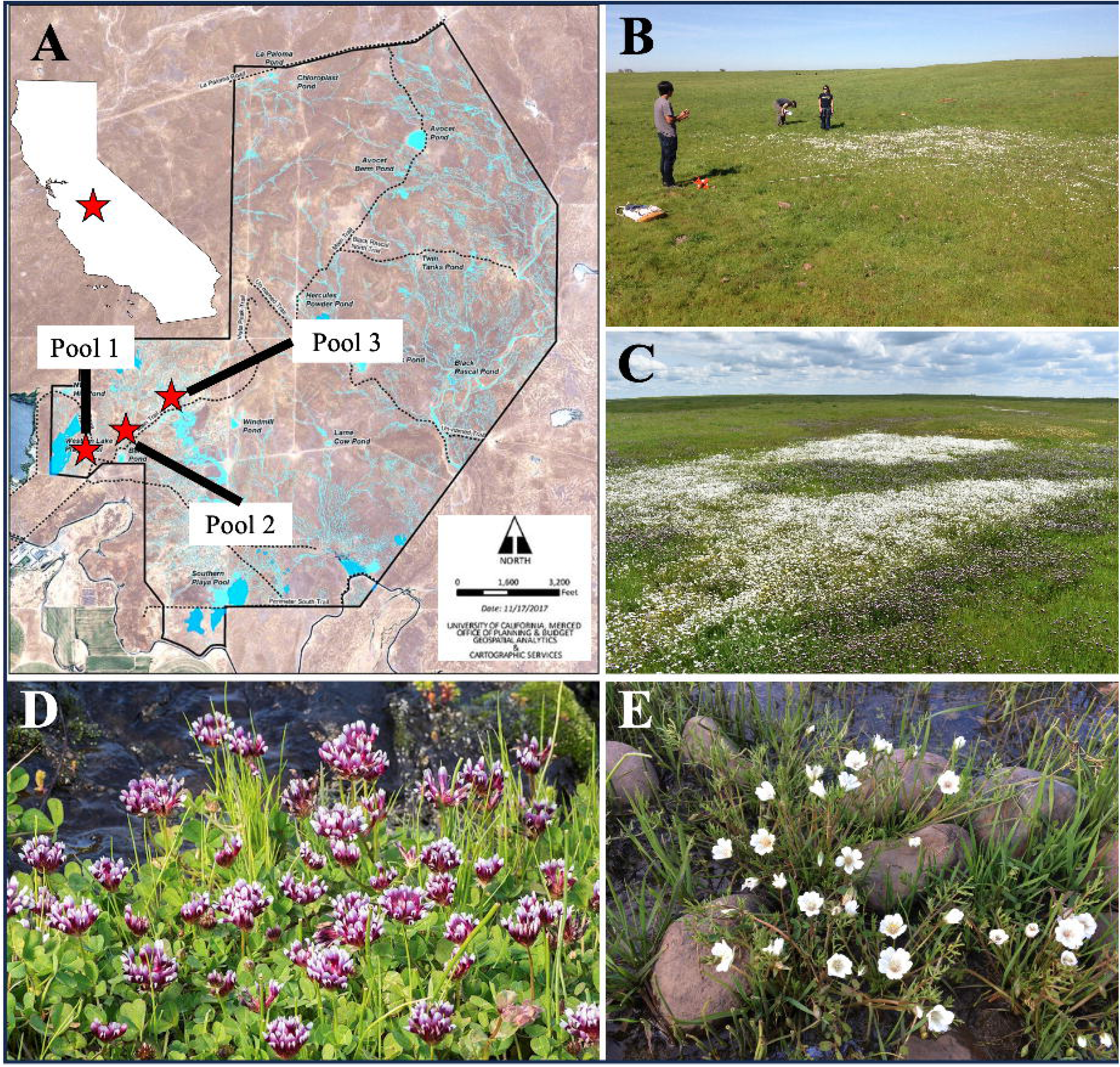
Summary of Merced climate from 2000-2022 and during the study period from 2016-2022. (A) Growing degree hours (top) and volume of rainfall in the early winter season (black line) and late winter season (grey line) with the study period marked by a dotted line. (B) Early winter average temperature (top) and late winter average temperature (bottom) metrics; daily maximum (red), daily mean (black), daily minimum (blue).

**Table 2:** Summary of the total growing degree hours, winter temperature, and volume of rainfall for the early and late winter seasons measured throughout the study period of 2016-2022. In addition, the day of first rainfall for each year is summarized in the last row.

| YEAR | 2016 | 2017 | 2018 | 2019 | 2020 | 2021 | 2022 |
| --- | --- | --- | --- | --- | --- | --- | --- |
| Early Winter GDH | 1190.00 | 1215.00 | 892.00 | 1152.00 | 1159.00 | 1142.00 | 1317.00 |
| Late Winter GDH | 1213.00 | 1187.00 | 1032.00 | 1046.00 | 1052.00 | 973.00 | 1007.00 |
| Early Winter Mean (C) | 11.13 | 10.85 | 10.39 | 10.75 | 11.16 | 10.82 | 11.40 |
| Late Winter Mean (C) | 10.44 | 9.89 | 9.35 | 8.96 | 9.68 | 9.52 | 9.23 |
| Early Winter Maximum (C) | 4.97 | 4.00 | 2.61 | 3.62 | 3.5 | 3.33 | 5.61 |
| Late Winter Minimum (C) | 4.77 | 4.43 | 3.07 | 3.32 | 2.86 | 2.58 | 1.59 |
| Early Winter Max (C) | 18.90 | 19.39 | 20.50 | 20.45 | 20.98 | 20.90 | 18.15 |
| Late Winter Max (C) | 16.84 | 16.12 | 16.55 | 15.70 | 17.90 | 17.46 | 18.48 |
| Early Winter Precipitation (mm) | 107 | 111.4 | 38.8 | 131.7 | 125.6 | 63.7 | 148.9 |
| Late Winter Precipitation (mm) | 255.6 | 325.6 | 94.6 | 226.4 | 79.2 | 142.1 | 34.6 |
| First Rainfall (Julian Date) | 274.00 | 289.00 | 292.00 | 275.00 | 329.00 | 311.00 | 294.00 |

Drier conditions were strongly correlated with higher swings in daily temperatures, which in turn lowered accumulated GDH. We found that CMI was negatively correlated with daily temperature range (p = 0.00257; ß = -0.25; R2 = 0.86, F = 31.04), suggesting that as conditions became moist, daily fluctuations of temperature were smaller. Furthermore, we found that when controlling for maximum daily temperature, larger diurnal temperature fluctuations resulted in lower GDH (p = 0.0041; ß = -91.58; R2 = 0.92, F = 21.74). We found that CMI was positively associated with GDH (p = 0.03821; ß = 52.20; R2 = 0.61, F = 7.81).

### 3.2 Phenological Patterns Over the Study Period

Meadowfoam and whitetip clover flowering start and end dates were earlier over time, whereas floral duration did not change. Meadowfoam’s day of first flower was significantly correlated with year (Figure 3), advancing at a rate of 4.7 days/year with a range of 50.7 days throughout the study period (2018: 89 days, 2022: 38.3 days). Similarly, whitetip clover’s day of first flower was negatively correlated with year (Figure 3), advancing at a rate of 5.6 days/year. The total change in the first flowering date from the start of the study (2016) to the end of the study (2022) was 22.4 days for meadowfoam and 30.4 days for whitetip clover. A significant negative correlation was detected for both species’ flowering end dates (Figure 3), with meadowfoam terminating flowering 4.9 days/year earlier and whitetip clover flowers ending 4.8 days/year earlier.

**Figure 3:**
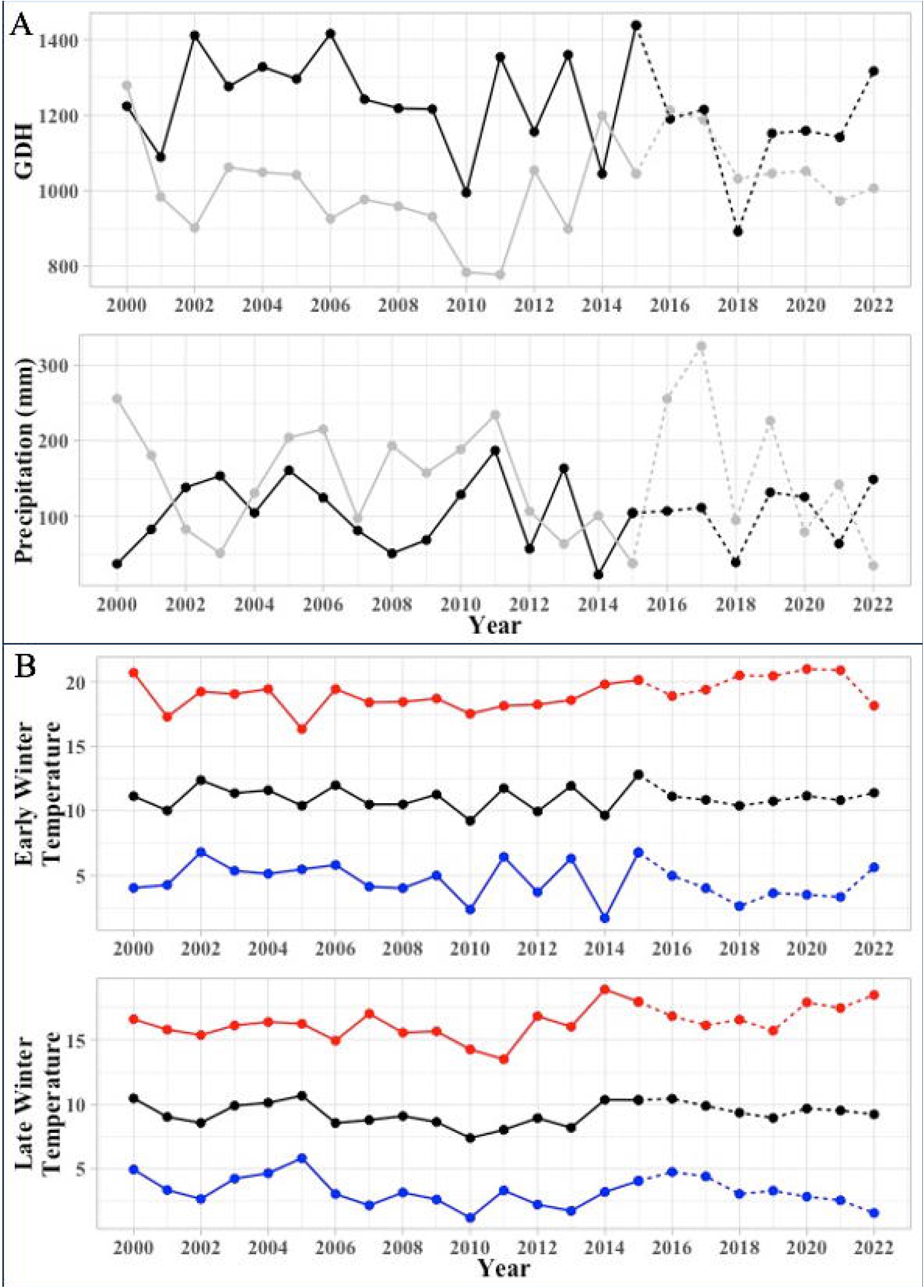
Linear regression of phenological variables (A) and density of phenophases (B) by year. Phenological variables were measured over 7 years, except for total and peak abundance of flowers which were measured from 2018 - 2022. Density variables were measured over a 4-year period from 2018-2021. Two average density measures were used: the first is the average number of plants, flowers, and fruits in all quadrats along a transect, and the second (If Present) is the average number of plants, flowers, and fruits in all quadrats where at least one plant was observed. Model summaries (R^2^, F-statistic, and p-value) of linear regressions are displayed at the top left corner of each graph. Each observation is an observation made at one of the three observation pools.

There was considerable variability in the flowering duration of plant populations. Whitetip clover populations were observed flowering continuously for 19 to 56 days, and meadowfoam populations flowered for a range of 32.7 days to 57.7 days. However, there was no distinct annual trend toward longer or shorter flowering periods for either species (Figure 3). The day to peak flowering – the date when the highest abundance of flowers was recorded for a single flowering season – significantly advanced at a rate of 5.7 days/year for meadowfoam and 5.1 days/year for whitetip clover (Figure 3).

### 3.3 Phenological Associations with Climate and Pool Characteristics

The phenology of both the generalist and specialist species showed a clear trend towards earlier flowering with warmer and drier winter conditions. Bloom start and end dates of meadowfoam and whitetip clover were negatively correlated with growing degree hours accumulated during the early winter (Figure 4). Conversely, growing degree hours accumulated during the late winter were not significantly associated with any phenological measurement (i.e., bloom start, bloom end, bloom duration, and peak flowering time) (Figure 4; Table 3).

**Figure 4:**
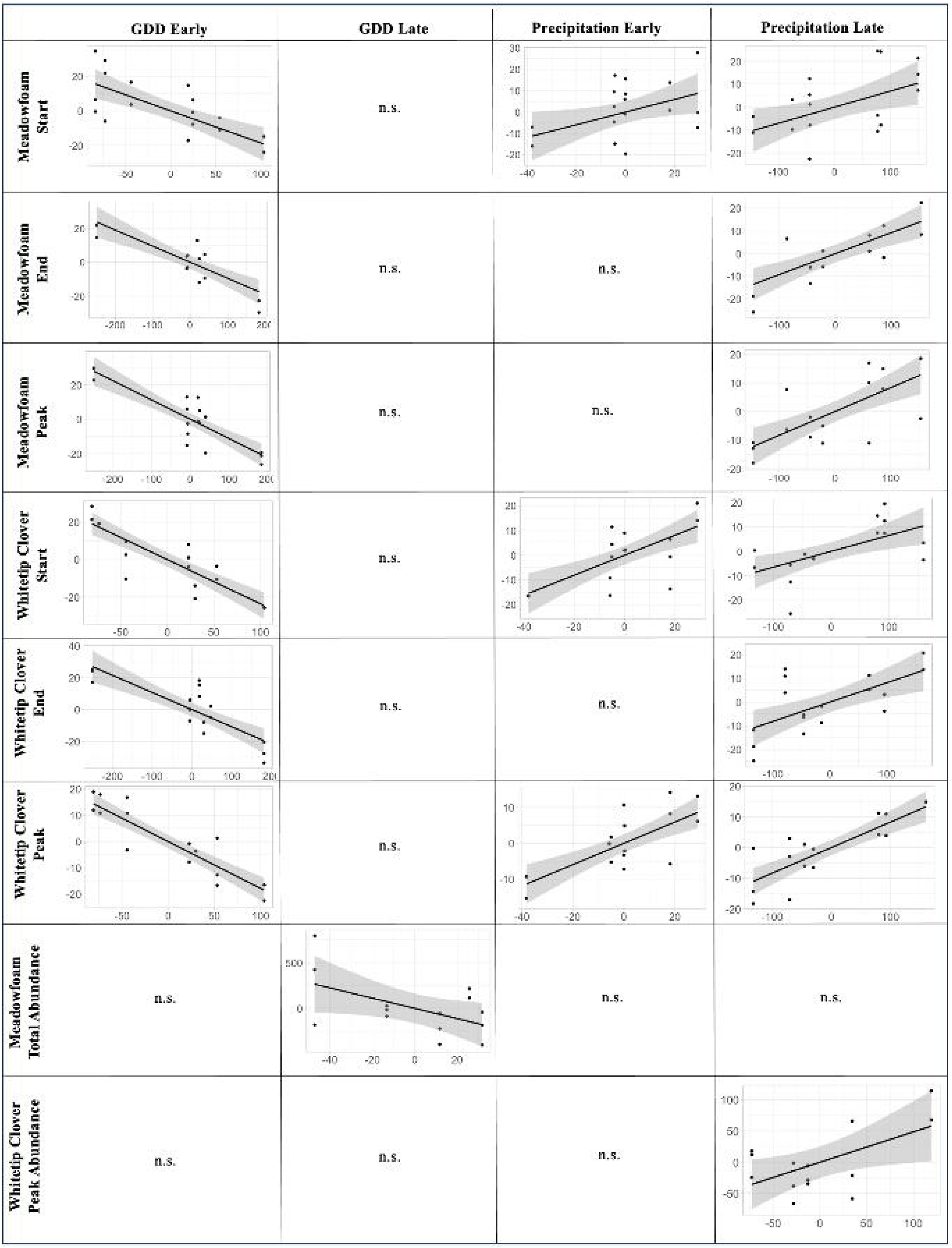
Added-variable plots for each phenological and population variable with at least one significant predictor from stepwise multiple linear modelling. Points are observations from one pool and one year. Best fit lines (solid line) and standard error (shaded region) are displayed for each significant association.

**Table 3.** Stepwise multiple linear regression model summaries of early and late winter GDH and precipitation as predictor variables of meadowfoam and whitetip clover phenological variables. Observations were made for 3 pools over 7 years. Pool 1 did not host any whitetip clover in 2019, reducing the number of observations to 20.

| Phenological Variable | Climate Predictor | Observations | Estimates | Std Error | P-value | R<sup>2</sup> |
| --- | --- | --- | --- | --- | --- | --- |
| Meadowfoam Start | Late Precipitation | 21 | 0.07 | 0.03 | 0.019 | 0.66 |
|  | Early Precipitation | 21 | 0.30 | 0.13 | 0.041 |  |
|  | Late GDH | 21 |  |  | ns |  |
|  | Early GDH | 21 | -0.19 | 0.04 | <0.0001 |  |
| Meadowfoam End | Late Precipitation | 21 | 0.09 | 0.02 | <0.0001 | 0.74 |
|  | Early Precipitation | 21 |  |  | ns |  |
|  | Late GDH | 21 |  |  | ns |  |
|  | Early GDH | 21 | -0.09 | 0.02 | <0.0001 |  |
| Meadowfoam Peak | Late Precipitation | 21 | 0.08 | 0.02 | <0.0001 | 0.80 |
|  | Early Precipitation | 21 |  |  | ns |  |
|  | Late GDH | 21 |  |  | ns |  |
|  | Early GDH | 21 | -0.11 | 0.01 | <0.0001 |  |
| Whitetip Clover Start | Late Precipitation | 20 | 0.06 | 0.02 | 0.006 | 0.85 |
|  | Early Precipitation | 20 | 0.40 | 0.10 | 0.001 |  |
|  | Late GDH | 20 |  |  | ns |  |
|  | Early GDH | 20 | -0.24 | 0.03 | <0.0001 |  |
| Whitetip Clover End | Late Precipitation | 20 | 0.08 | 0.02 | 0.002 | 0.74 |
|  | Early Precipitation | 20 |  |  | ns |  |
|  | Late GDH | 20 |  |  | ns |  |
|  | Early GDH | 20 | -0.11 | 0.02 | <0.0001 |  |
| Whitetip Clover Peak | Late Precipitation | 20 | 0.08 | 0.01 | <0.0001 | 0.90 |
|  | Early Precipitation | 20 | 0.30 | 0.06 | <0.0001 |  |
|  | Late GDH | 20 |  |  | ns |  |
|  | Early GDH | 20 | -0.18 | 0.02 | <0.0001 |  |

Higher precipitation was associated with delayed phenological patterns for both species. Early winter precipitation was positively associated with meadowfoam and whitetip clover start dates (Figure 4). Late winter precipitation was positively correlated with meadowfoam and whitetip clover start dates, end dates, and peak dates (Figure 4).

We found that all phenological variables were positively associated with late winter mean temperature and negatively associated with early winter mean temperature (Supplemental Table 1). Furthermore, we found that both focal species advanced their flowering start, termination, and peak abundance dates in response to warmer early winter minimum temperatures. In contrast, warmer late winter minimum temperatures resulted in delayed flowering termination and peak abundance dates (Supplemental Table 1). When maximum temperature was higher in the late winter, significant advances of meadowfoam and whitetip clover start dates were observed.

The CMI of late winter was positively associated with the start, end, and peak bloom dates of both focal species (Supplemental Table 1).

Despite no climatic association with floral duration, local microtopographic features were related to a strong phenological response in the vernal pool specialist, meadowfoam. The number of days meadowfoam plants flowered was longer near the pool center (Table 4; Supplemental Figure 3). This relationship is held for each collection pool and year observed. The floral duration of whitetip clover did not respond to its location in the pool (Table 4).

**Table 4.** Linear regression model summaries of meadowfoam and whitetip clover phenological variables with distance from edge as a predictor variable. Observations are quadrats with at least one meadowfoam or whitetip clover plant collected across three pools and 7 years.

| Phenological Variable | Observations | Estimate | Std Error | P-value | R<sup>2</sup> |
| --- | --- | --- | --- | --- | --- |
| Meadowfoam Start | 1085 | -0.30 | 0.22 | 0.169 | 0.00 |
| Meadowfoam End | 1085 | 1.07 | 0.16 | <0.0001 | 0.04 |
| Meadowfoam Length | 1085 | 1.37 | 0.18 | <0.0001 | 0.05 |
| Whitetip Clover Start | 946 | 0.34 | 0.22 | 0.122 | 0.00 |
| Whitetip Clover End | 946 | 0.37 | 0.17 | 0.033 | 0.01 |
| Whitetip Clover Length | 946 | 0.02 | 0.14 | 0.885 | 0.00 |

### 3.3 Phenology and Climate Effects on Populations

The rate at which meadowfoam and whitetip clover were found within pools was strongly associated with both climatic and phenological variability. Average occupancy rates of meadowfoam and whitetip clover, both with and without standardizing by the number of quadrats per pool, were positively associated with winter precipitation and negatively associated with winter GDH (Table 5). Furthermore, we found that the earlier flowers bloomed, the higher the occupancy rates of meadowfoam and whitetip clover were. Delayed termination dates of whitetip clover were associated with greater occupancy rates.

**Table 5.**
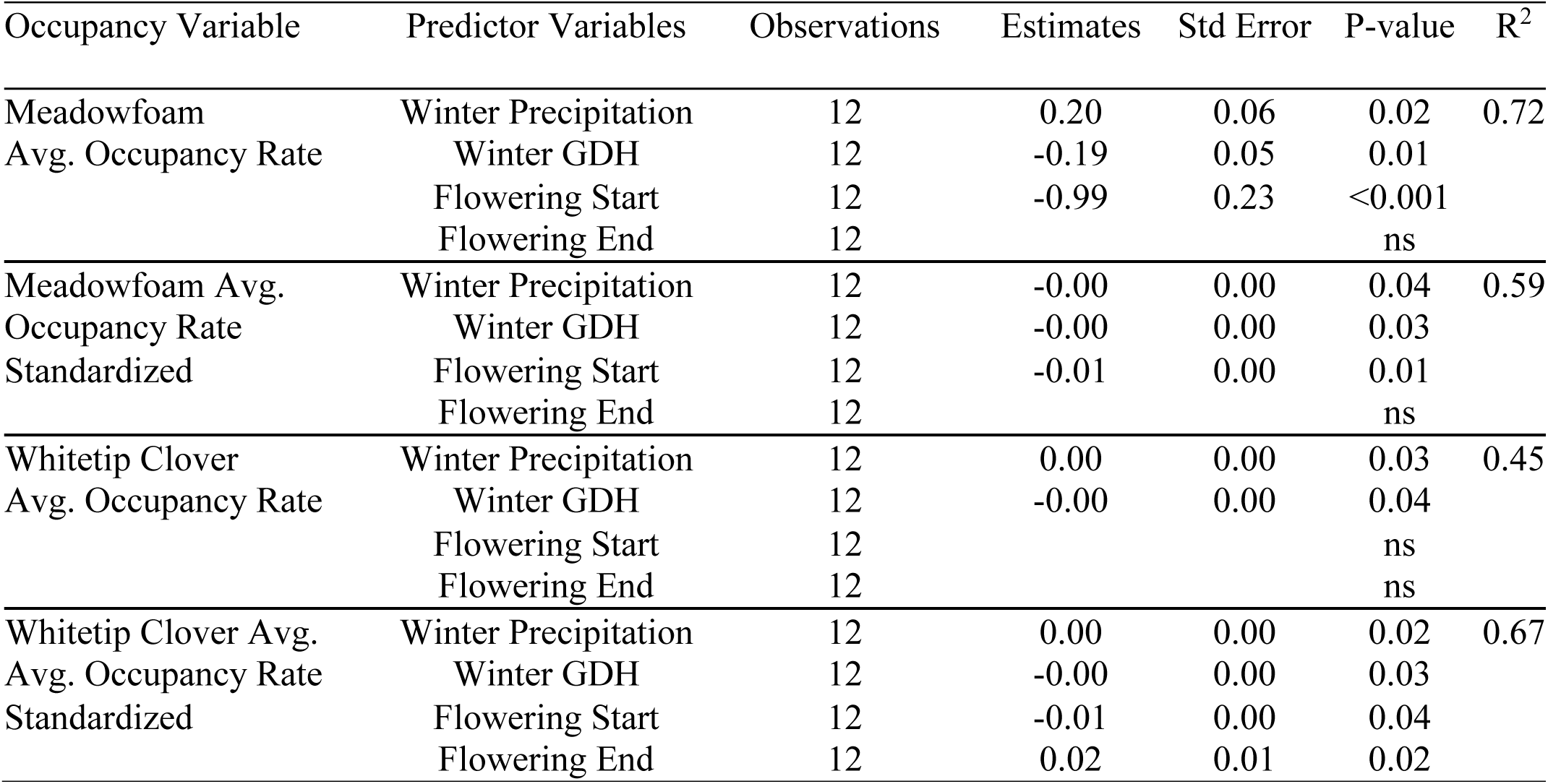
Stepwise multiple regression of meadowfoam and whitetip clover average occupancy rate across all quadrats within a pool and average occupancy rate standardized by the number of quadrats within a pool. Observations are 3 pools across 4 years.

Divergent patterns of plant density and fecundity were observed over the study period. From 2016 to 2022, the density of meadowfoam and whitetip clover decreased while floral and fruit density increased (Figure 3).

Density of plants, flowers, and fruits was significantly associated with climate and phenology, though the responses differed between species. Plant and flower plot density of meadowfoam were positively associated with greater precipitation (Supplemental Table 2). The plot density of whitetip clover plants responded similarly to greater precipitation, though whitetip clover fruit density was negatively correlated with precipitation, and flowers were unaffected. Furthermore, greater GDH was correlated with reduced whitetip clover plant density and positively correlated with whitetip clover fruit density. The floral density of whitetip clover was negatively correlated with start dates, and neither species exhibited changes in density due to termination dates.

## DISCUSSION

Ephemeral wetlands exhibit high interannual and seasonal variation in inundation, making multi-year studies crucial for understanding community responses to a range of climatic conditions (Pätzig et al., 2020). In this study, we examined vernal pools over a span of 7 years, encompassing climate years with both relatively high and low precipitation compared to the past two decades, as well as a wide variation in annual accumulation of growing degree hours. Thus, the current dataset’s time period is advantageous for parameterizing the response of natural plant floral phenology to climate.

We found that temperature, precipitation, and the CMI were highly correlated with the phenological timing of meadowfoam and whitetip clover in vernal pool environments. The climate conditions of early and late winter seasons exerted distinct effects on plant phenology, underscoring the importance of capturing a broad seasonal variation that influences germination, vegetative growth, and transitions to reproduction. These findings highlight both the inherent variability of vernal pool plant phenology and the significant impact that warmer and drier conditions have on the flowering times of widespread and endemic species. To our knowledge, this study represents the first examination of climatic drivers of vernal pool plant phenology in the field and offers insights into how phenology may be affected in these vulnerable ecosystems by future climate change.

### 4.1 Phenological Patterns and Associations with Climate

Meadowfoam, a vernal pool specialist, and whitetip clover, a widespread vernal pool associate, both initiated, terminated, and reached peak flowering earlier under warmer and drier conditions in natural vernal pools. Warming promotes earlier germination and growth of vernal pool species (Bliss & Zedler 1998), and we found that floral phenology also advanced in response to higher annual temperatures. Laboratory studies of meadowfoam under various temperature conditions revealed faster growth and seed development at higher temperatures, indicating a natural sensitivity to warming (Franz 1989). While no previous study has examined the effect of warming on whitetip clover, evidence of phenological responsiveness to temperature exists in other clover species such as *Trifolium repens* (Murray et al., 1999), *Trifolium pratense* (Hulme 2011), and *Trifolium andersonii* (Kopp & Cleland 2015). These patterns align with those observed in other spring wildflower species that exhibit advanced flowering in response to hotter seasonal conditions (Parmesan & Yohe 2003; Cook et al., 2012; Pearson 2019).

The specialist and generalist vernal pool plant species exhibited similar phenological responses to precipitation, temperature, and drought. Overall, phenological variables of both species showed surprisingly similar responses to climate variation (Table 3), except for early winter precipitation, which affected whitetip clover start dates more than meadowfoam start dates. The slopes of response for both species’ start dates in response to early winter growing degree hours (GDH) were twice those for end dates, indicating a stronger relationship between early winter growing seasons and floral initiation than between early winter growing seasons and floral termination. Nearly all phenological metrics of both species were associated with mean, minimum, and maximum temperatures (Supplemental Table 1), suggesting that temperature is a good predictor for vernal pool species phenology. A positive association of late winter CMI with phenology, where higher CMI indicates moister and less drought-like conditions, suggests delayed phenology when more moisture is available. The response rate of each respective phenological variable was similar for both species, indicating that vernal pool specialists and generalist species adjust phenology similarly in response to drought conditions.

Our findings indicate that early growth strongly influences floral phenology later in the life cycle of meadowfoam and whitetip clover, underscoring the importance of monitoring warming trends in both early and late winter. Vernal pool plants germinate and grow slowly during the fall and early winter (Keeley & Zedler 1998), with most of the growth typically observed in spring. Meadowfoam and whitetip clover phenology responded strongly to early winter temperatures. In a study of constructed vernal pools at Travis Air Force Base, California, focusing on 11 native and non-native plant species, Javornik & Collinge (2016) found significant associations of late winter Growing Degree Days (GDD) with plant emergence during 2002-2012, but no association with early winter GDDfor native or non-native species. . Recent evidence suggests that early growth can play a pivotal role in later floral phenology of the vernal pool associate plant, *Lasthenia californica*. Olliff-Yang & Ackerly (2021) found that *L. californica* populations experimentally seeded in serpentine grasslands in November reached the size threshold for inducing flowering significantly earlier than those seeded in January and March. These results imply that important development occurs between November and January, the period considered early winter in this study, that can impart a significant effect on spring phenology.

The delayed phenological patterns observed in 2018, which was abnormally cold and dry during both early and late winter seasons, highlights the importance of considering seasonal conditions that influence all stages of a plant’s development. Despite the general trend that dry winter conditions accelerate spring flowering, colder temperatures in 2018 may have limited development during the early winter. Accumulated GDH in 2018 was significantly lower than in all other recorded climate years during the study. We hypothesize that poor vegetative growth resulting from low precipitation and cold, biologically restrictive temperatures in 2018 delayed spring flowering contrary to phenological expectations under dry conditions. Such delayed flowering, because of delayed germination and vegetative growth, aligns with findings by Olliff-Yang & Ackerly (2021), although their study occurred in a different habitat type. Monitoring early season vegetative performance would be necessary to confirm our interpretations.

First flowering date may be more responsive to climate variation than floral duration. Neither meadowfoam nor whitetip clover showed a significant effect of precipitation and temperature on floral duration. Our findings align with those of *Anemone nemorosa*, an herbaceous wildflower, which exhibited a significant advancement of first flowering dates in response to rising temperatures but no significant shortening of the flowering period in temperate forests (Pulchalka et al., 2022). Similarly, a community-scale study on Guernsey island in the English Channel, involving 232 plant species monitored over 27 years, found that 55% had significantly advanced first flowering dates, while only 19% exhibited a shorter floral duration (Brock et al., 2014). In contrast to our findings, simulated warming shortened the flowering duration in *Arabidopsis helleri* ssp. *gemmifera* (Nagahama et al., 2018) and *Cardamine hirsuta* (Cao et al., 2016) in controlled growth chambers. Shorter flowering durations resulted from delayed floral initiation and advanced floral termination for *A. helleri* subsp. *gemmifera* and *C. hirsuta*, respectively. Future research will be necessary to connect molecular pathways with observed trends of floral duration in the field to resolve disparities between these biological scales.

Our findings indicate that inter- and intra-annual precipitation variability drives shifts in floral phenology of vernal pool species, drawing parallels with some plant habitats while differing from others. Inundation plays a crucial role in wetland plant establishment, germination, and survival (Diel 2005), structuring spatial and temporal patterns of community composition in vernal pool habitats (Bauder 2005; Emery et al., 2009; Javornick & Collinge 2016). Fall and winter precipitation are primary water sources in vernal pools (Jokerst 1990), and interannual rainfall variability promotes changes in community composition and phenology (Javornick & Collinge 2016; Martin & Lathrop 1986; Collinge et al., 2003). Higher soil moisture and delayed soil desiccation dates are generally associated with later flowering among vernal pool associates within genera such as *Lasthenia*, *Downingia*, and *Pogogyne* (Martin & Lathrop 1986; Schiller et al., 2000; Collinge et al., 2003). Similarly, we observed delayed start, end, and peak dates of flowering in response to higher accumulated rainfall for meadowfoam and whitetip clover. A strong relationship between phenology and hydrology is also found among obligate aquatic plants in ponds and lakes (Calero et al., 2015), where water depth, rather than soil desiccation timing, drives life history transitions of submerged and emergent vegetation. Unlike aquatic plants, it remains uncertain whether changing precipitation patterns cause significant phenological shifts in terrestrial species found in temperate (Sparks et al., 2006), alpine (Hart et al., 2014), Mediterranean (Gordo & Sanz 2005), and coastal regions (Cleland et al., 2006). However, current knowledge suggests that precipitation plays a clearer role in modulating floral phenology in tropical (Zalamea et al., 2011), subtropical (Peñuelas et al., 2004; Pearson 2019), and arid regions (Crimmins et al., 2013).

Vernal pool species are present as segregated communities along the pool gradient according to specific adaptations (Schlising & Sander 1982; Zedler 1984; Hendrickson 2024). This spatial and temporal segregation in vernal pools arises from edaphic and hydrologic differences along the gradual slope of the pool (Holland & Jain 1984; Zedler 1984). In this study, we found temporal niche partitioning along the inundation gradient, as the onset and peak date of meadowfoam flowering occurred, on average, 10 days earlier than whitetip clover. Temporal niche segregation is a critical phenomenon for species coexistence (Wiens et al., 1977), particularly in environments with limited space such as vernal pools. Although we did not find that climate variation influences floral duration, we observed that the topography of vernal pools, along with inundation length and soil moisture, affects the floral duration of the vernal pool specialist more than the generalist species. Our observations indicate that meadowfoam phenology is more sensitive to moisture and clay content compared to whitetip clover. This is supported by findings that meadowfoam floral duration and density were greater nearer deeper regions of the pool with higher soil moisture. In contrast, whitetip clover showed no phenological response to its location within the pool, with equal flowering frequency observed at the edge and bottom zones. These results suggest that while climate influences both species similarly, the phenological sensitivity to soil moisture, promoted by the unique topography of vernal pools, is greater for the specialist species than the generalist species.

### 4.2 Implications for Climate Change

Our results highlight the potential for a dramatic advancement in mean flowering times within a relatively short period, consistent with climate change predictions. Over the past 30 years, the climate of the California Central Valley has been trending towards warmer winter and spring seasons (Dettinger & Cayan 1995; He et al., 2017). These shifts have led to significant advances in flowering times among many annual species (Lesica & Kittelson 2010; Abu-Asab et al., 2001; Primack et al., 2004). Hydrological models of future climates predict shorter inundation periods for California’s vernal pools due to increased warming and evaporation rates, potentially endangering vernal pool specialists adapted to longer inundation periods (Montrone et al., 2019). In our study, we found that both facultative and obligate vernal pool species exhibit profound phenological responsiveness to precipitation and temperature by advancing flowering times under drier and warmer climatic conditions. This suggests that projected climate changes in the Central Valley will likely drive predictable advancements in vernal pool plant phenology. To underscore the potential change in floral phenology, we observed that during the recent three-year megadrought in California, meadowfoam initiated flowering in February, a month earlier than reported by the Jepson eflora herbarium and a month earlier than all field observations within the Calflora database. These phenological patterns suggest that if drought conditions become more frequent, as projected (Berg & Hall 2015; He & Gautam 2016), mean flowering onset is likely to remain advanced by a month or more, with unknown ecological consequences.

Long-term phenological observations in threatened and vulnerable ecosystems are critical for understanding and responding to global change threats. Our observations indicate that both vernal pool specialists and widespread spring wildflowers are sensitive and susceptible to potential climate changes in the Central Valley of California. However, plant species from warmer and more variable climates, such as Mediterranean climates, may exhibit higher levels of phenotypic plasticity in various morphological and phenological traits (Kreyling et al., 2019) that could help mitigate or buffer populations somewhat from future climate stresses. We noted high interannual variation in phenology, suggesting the capacity for phenotypic plasticity in meadowfoam and whitetip clover, although further research is needed to confirm significant gene-environment effects. Importantly, we found that a shifted flowering schedule was not accompanied by a reduction in fruiting abundance or plant density for either species. Future research should examine whether pollinators can track changes in vernal pool plant phenology, providing insights into community-level consequences of future climate variability.

## Acknowledgements

We thank many student volunteers and interns, including Aime Arreola, Azul Carrillo, Madeline Castro, Tiffany Chang, Andrea Diaz Cruz, Marcel Dube, Kevin Felix, Arlene Garcia, Andrew Ho, Daniel Hiebert, Hedaq Ibrahim, Martha Kandziora, Eloise Moa, Dulce Monje Alcazar,Iris Montes, Leslie Paredes, Avalon Patton, Zara Perez-Ochoa, Antonio Pina, Mimi Pomephimkham, Sean Quarnstrom, Danny Rivas, Valeria Rivera, Alexander Rogado, Elizabeth Souvannarath, and Ash Valdez. Special thanks to students Robert Martin, Julia DePass, and Jenna Heckel for helping to establish this project as part of the Yosemite Leadership Program Capstone Project. We thank past directors of the UC Merced Vernal Pool and Grasslands Reserve, Monique Kolster and Chris Swarth, and current Director Joy Baccei, for use of the reserve and for their great support, including use of vehicles, supplies, and databases. We thank Jacob Croasdale, Jack Cronin, and Daniel Toews, Susan Mazer, Teamrat Ghezzehei, and Jessica Blois for assistance and advice during this project. Support for this project was provided to JPS from a grant from the US Fish and Wildlife Service and Bureau of Reclamation’s Central Valley Project Conservation Program (CVPCP) and Central Valley Project Improvement Act Habitat Restoration Program (HRP) (CESU—R17AC00044).

## Author Contributions

BTH : Conceptualization, Methodology, Validation, Formal Analysis, Investigation, Data Curation, Writing Original Draft, Writing Review & Editing, Visualization, Project Administration

AH : Conceptualization and Investigation

RM : Conceptualization and Investigation

JS: Conceptualization, Methodology, Validation, Investigation, Writing Review and Editing, Supervision, Project Administration, and Funding Acquisition.

### Data Availability

The data used to produce results for this study are available at figshare (10.6084/m9.figshare.25065977.v1) and code is available on github (https://github.com/Brandon-Thomas-Hendrickson/Meadowfoam_WhitetipClover_Phenology.git).

