## Supplemental Tables for "Climatic determinants of plant phenology in vernal pool habitats"

Supplemental Table 1: Stepwise multiple linear regression model summaries of (A) mean temperature, (B) minimum temperature, and (C) maximum temperature with precipitation as predictor variables of meadowfoam and whitetip clover phenological variables. (D) Early and late winter climate moisture index (CMI) with precipitation as predictor variables f meadowfoam and whitetip clover phenology.

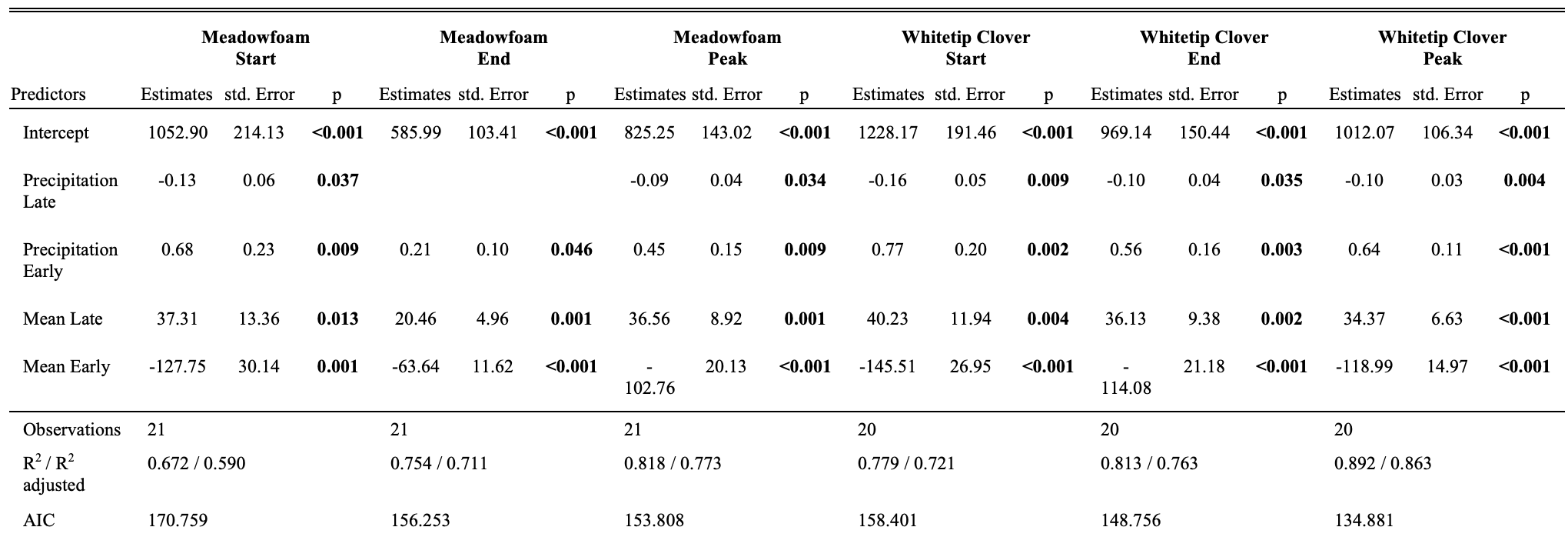

**A**

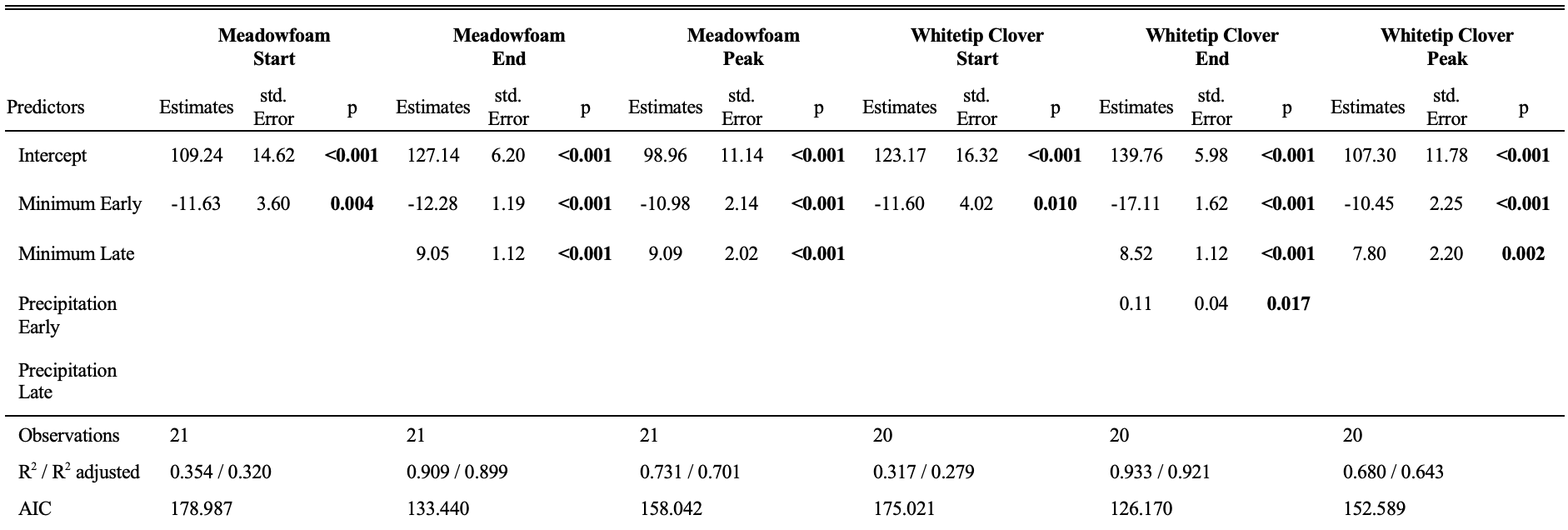

**B**

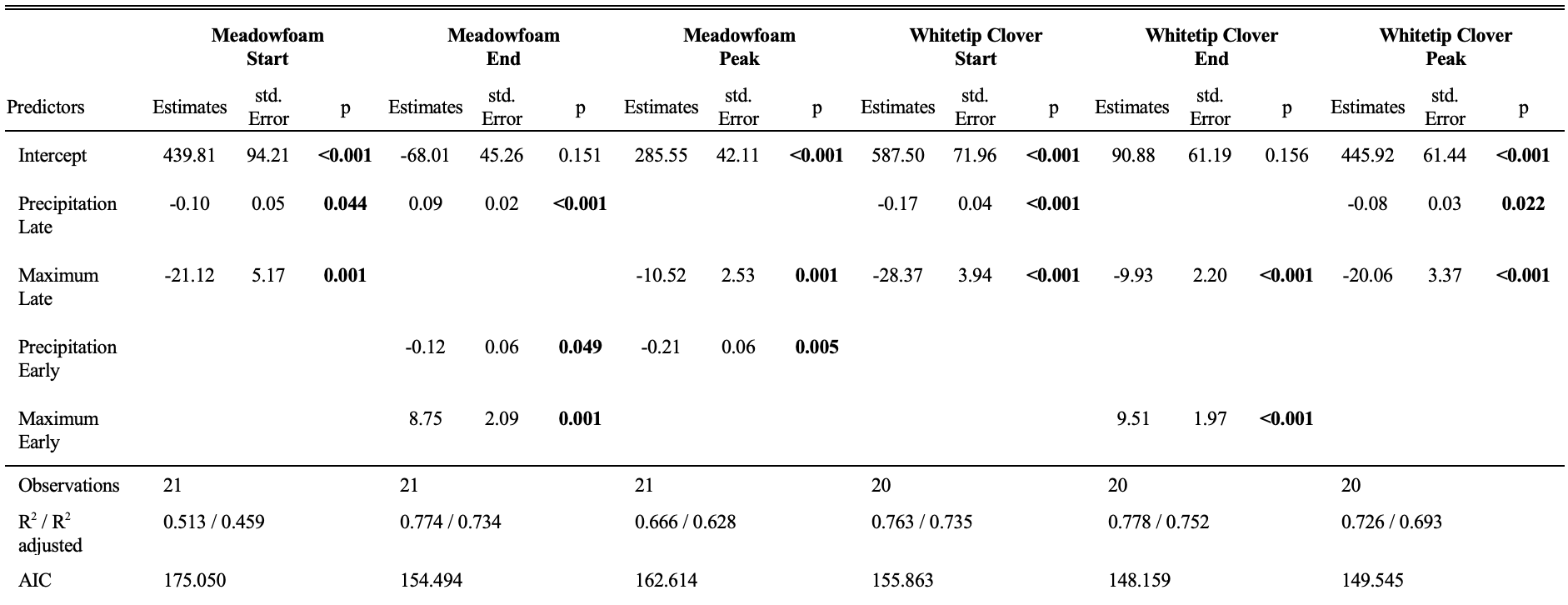

**C**

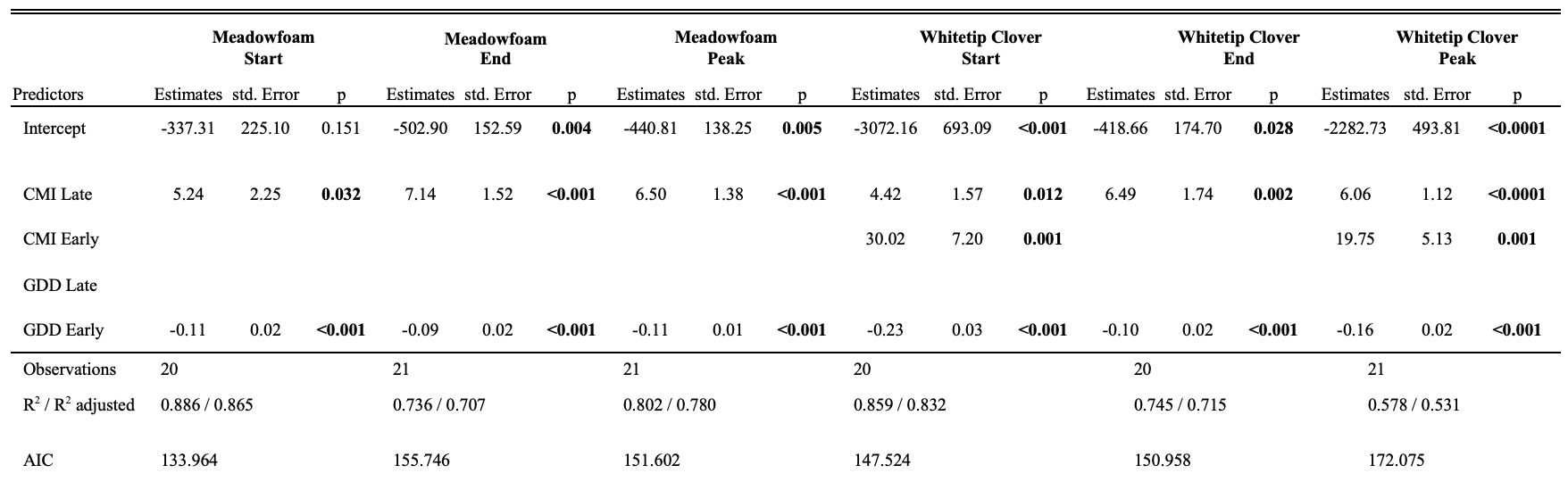

**D**

Supplemental Table 2: Stepwise multiple regression of meadowfoam and whitetip clover plant, flower, and seed density of quadrats with at least one plant observed by winter precipitation, winter GDH, flowering start date, and flowering end date as predictor variables. Observations are 3 pools across 4 years. Associations reported had at least one significant predictor variable, whereas meadowfoam seed density was not significantly associated with any predictor. Furthermore, no density variable was significantly associated with any predictor variables when all quadrats in a pool was examined.

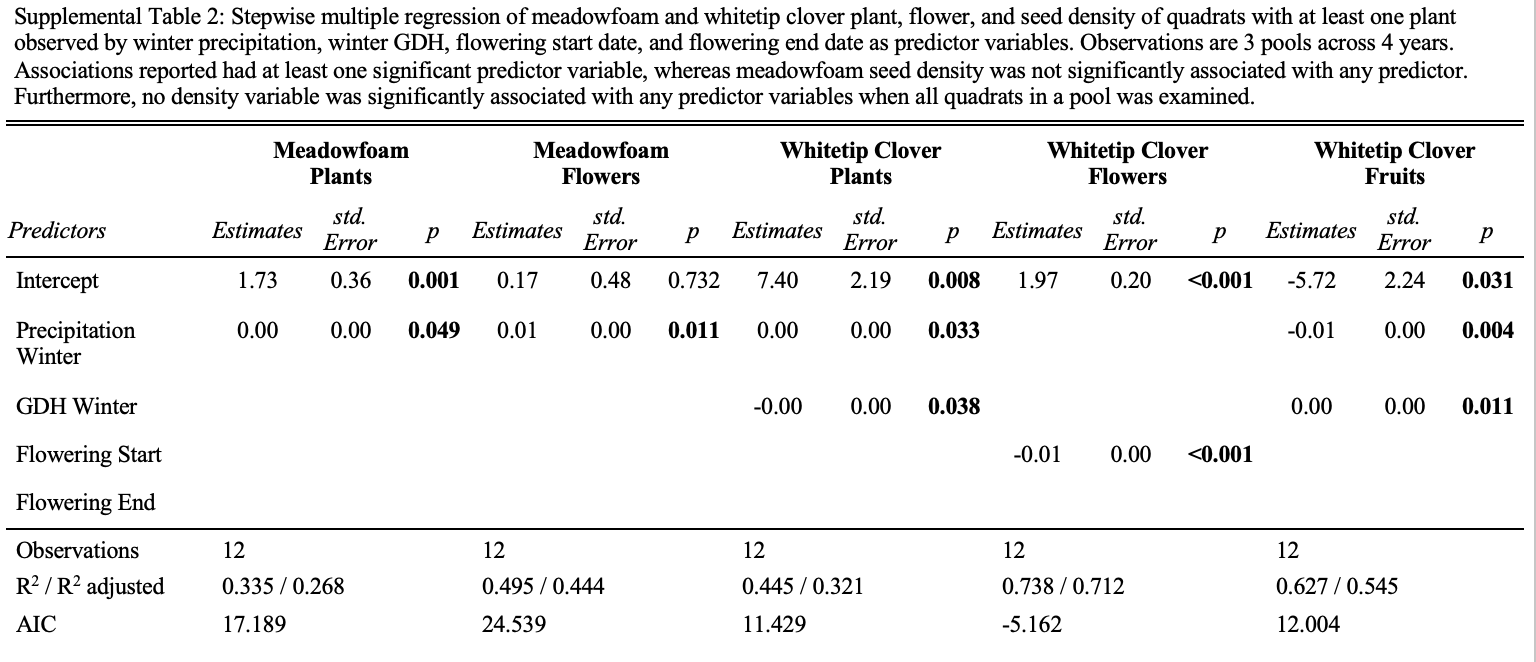
