## Supplemental Figure Legends for "Climatic determinants of plant phenology in vernal pool habitats"

Supplemental Figure 1: Early winter precipitation (A) and late winter precipitation (B) boxplots of five CIMIS stations. The control station, Merced, was contrasted with the four stations ranging from 16km to 160km away from Merced. Dunnett’s model p-values are displayed above each station that was contrasted with the control.

Supplemental Figure 2: Boxplots of (A) meadowfoam and (B) whitetip clover phenology and population measures for three observation pools (x-axis) recorded from 2016 – 2022.

Supplemental Figure 3: Meadowfoam and whitetip clover start (blue) and end (red) dates along the transect of pool 1 (top), pool 2 (middle), and pool 3(bottom). The shaded region is the length of flowering time, which is then depicted on the right-hand panel of each.
