## Supplementary figures and images for "Climatic determinants of plant phenology in vernal pool habitats"

### Supplemental Figure 1

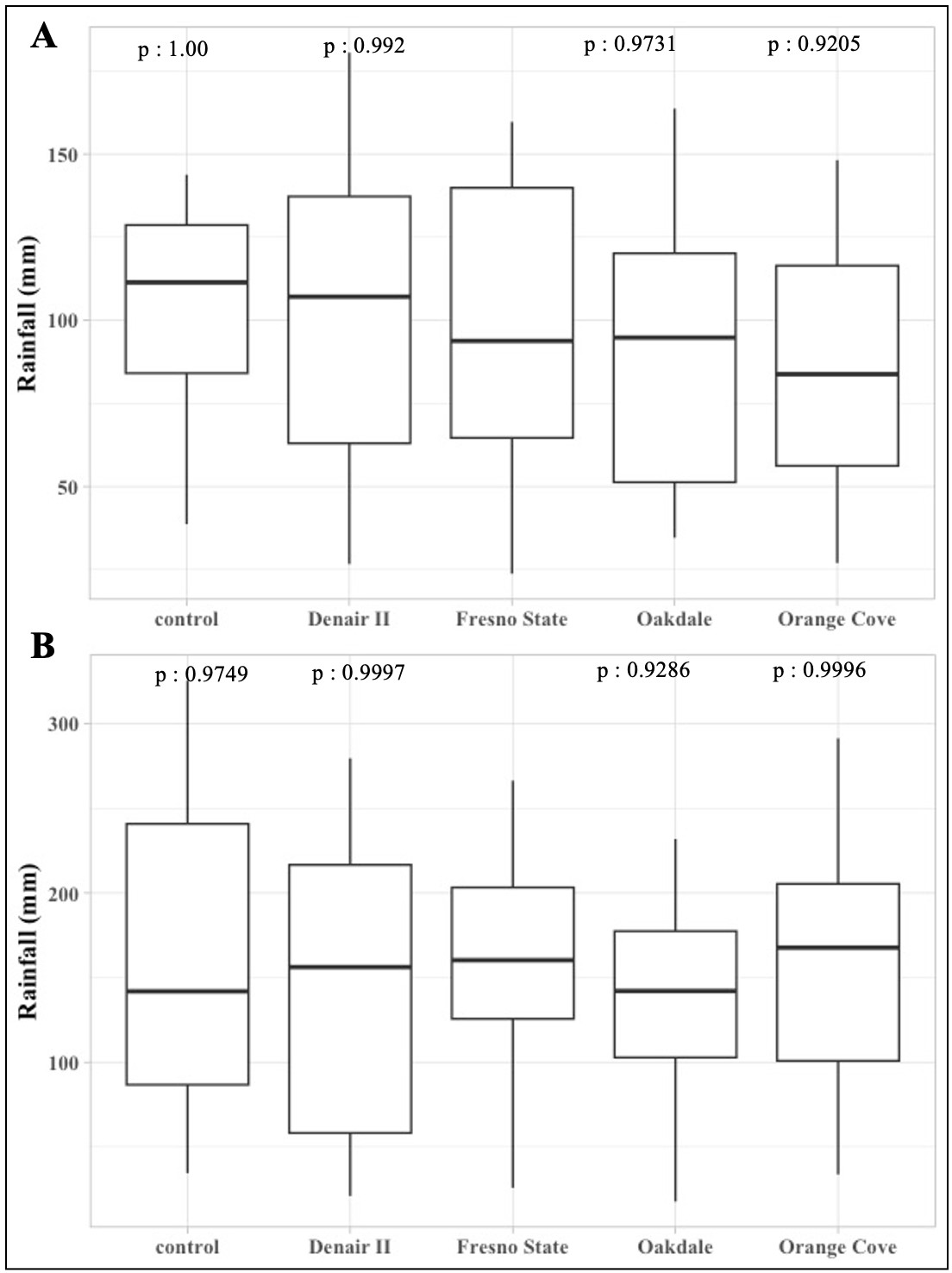

### Supplemental Figure 2

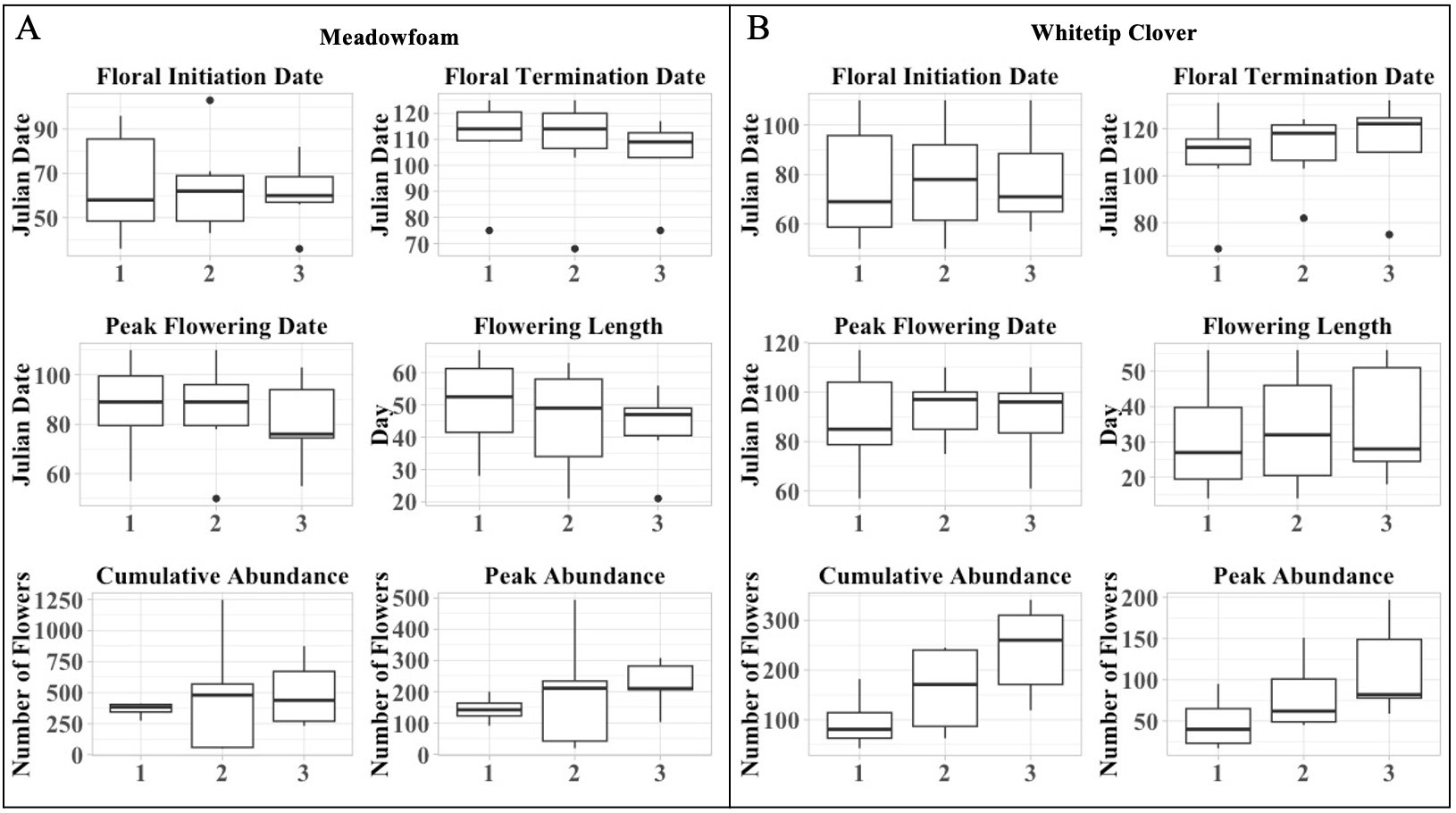

### Supplemental Figure 3

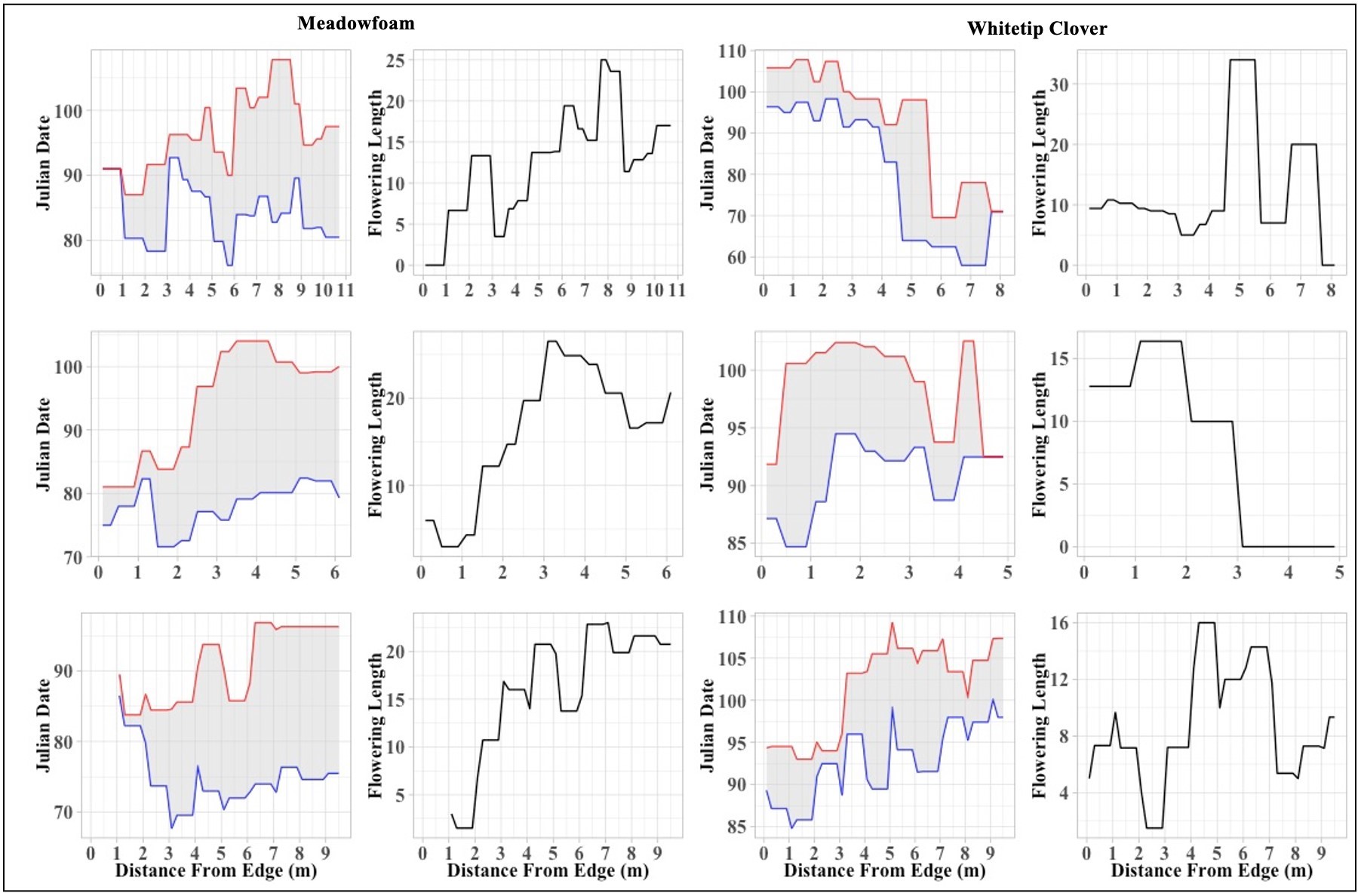
